# Functional Sexual Dimorphism in Human Nociceptors

**DOI:** 10.1101/2023.06.14.545010

**Authors:** Harrison Stratton, Mahdi Dolatyari, Nicolas Dumaire, Aubin Moutal, Andre Ghetti, Tamara Cotta, Stefanie Mitchell, Xu Yue, Edita Navratilova, Frank Porreca

## Abstract

The recent demonstration of differences in transcript expression in human post-mortem sensory neurons suggests the possibility of sexually dimorphic pain mechanisms. To date, however, the concept of “male” and “female” nociceptors has not been demonstrated at a functional level. We now report sensitization of female, but not male, human nociceptors by prolactin revealing a female-selective mechanism that can be exploited to improve the treatment of pain in women.

## Main text

Women are overrepresented in many pain conditions^1-3^, especially those that occur without evidence of tissue pathology such as migraine, irritable bowel syndrome, temporomandibular disorder, painful bladder syndrome and fibromyalgia^4-8^. Additionally, many pain conditions such as endometriosis, dysmenorrhea and vulvodynia are female specific^9-11^. An emerging concept that could be relevant to female prevalent or specific pains is that nociceptors, afferent neurons that transmit potentially tissue damaging stimuli from the periphery to the central nervous system, may be sexually dimorphic^12, 13^. Differences in transcript, protein and function have emerged from analysis of dorsal root ganglion (DRG) neurons of male and female rodents^14, 15^. Differences in transcripts of DRG neurons from male and female human donors have also been reported supporting that there may be male and female nociceptors^16^. To date, however, there are no reports of functional differences between nociceptors from male and female human donors. While data from animal DRG neurons correlate well with emerging information from human cells, important differences have also been reported. This includes the failure to replicate in humans the broad classification of DRG neurons as “peptidergic” and “non-peptidergic” derived in animal studies^17^. Confirmation of functional sex differences in human nociceptors would support the novel possibility of implementation of precision medicine by consideration of patient sex in choice of pain therapies.

Peripheral nociceptor sensitization can promote increased pain by decreasing activation thresholds leading to inappropriate engagement of high threshold afferent transmission pathways by low intensity stimuli^18^. Many substances sensitize rodent nociceptors, but few reports reveal preferential sensitization between sexes. One sensitizing agent is prolactin, a neurohormone released by the pituitary and influenced by estrogen and stress^19^. Women and female animals have higher circulating prolactin levels and women demonstrate stronger stress responses than men^20^. Prolactin is additionally produced locally by a number of different cell types^21^ and both pituitary and extra-pituitary prolactin selectively sensitizes female nociceptors in rodents^14^. The female selectivity of prolactin in rodent nociceptor sensitization has been confirmed in both cellular and in vivo studies^22^. Confirmation of female-selective human nociceptor sensitization would increase understanding of female prevalent, and female specific, pain disorders and support development of female-specific pain therapies.

We explored the possibility that prolactin might differentially sensitize DRG neurons recovered from male and female human donors to determine (a) if functional sexual dimorphism occurs in human nociceptors and (b) whether targeting prolactin could represent a therapeutic strategy with high likelihood of translation across species. We identified the concentration at which exogenous mouse prolactin (mPRL) produced sensitization of mouse DRG cells using patch clamp electrophysiology. We crossed mice that express Cre recombinase under the control of the prolactin receptor promoter (Prlr^Cre/+^) with a GFP reporter line Ai6 (see methods). We cultured lumbar DRG neurons from adult female Prlr^Cre/+^;Ai6 mice (**Fig. 1A**) and visualized Prlr expressing GFP positive neurons (**Fig. 1B**). These neurons were incubated overnight with graded concentrations recombinant mPRL and neuronal excitability was measured the following day with action potential firing frequency as a functional endpoint; mPRL increased excitability of cells with an EC50 of ∼29 nM (**Fig. 1C**). From the concentration-response curve, incubation with 50 nM PRL was established as a saturating concentration and used in further studies (**Fig. 1D**).

**Figure 1.**
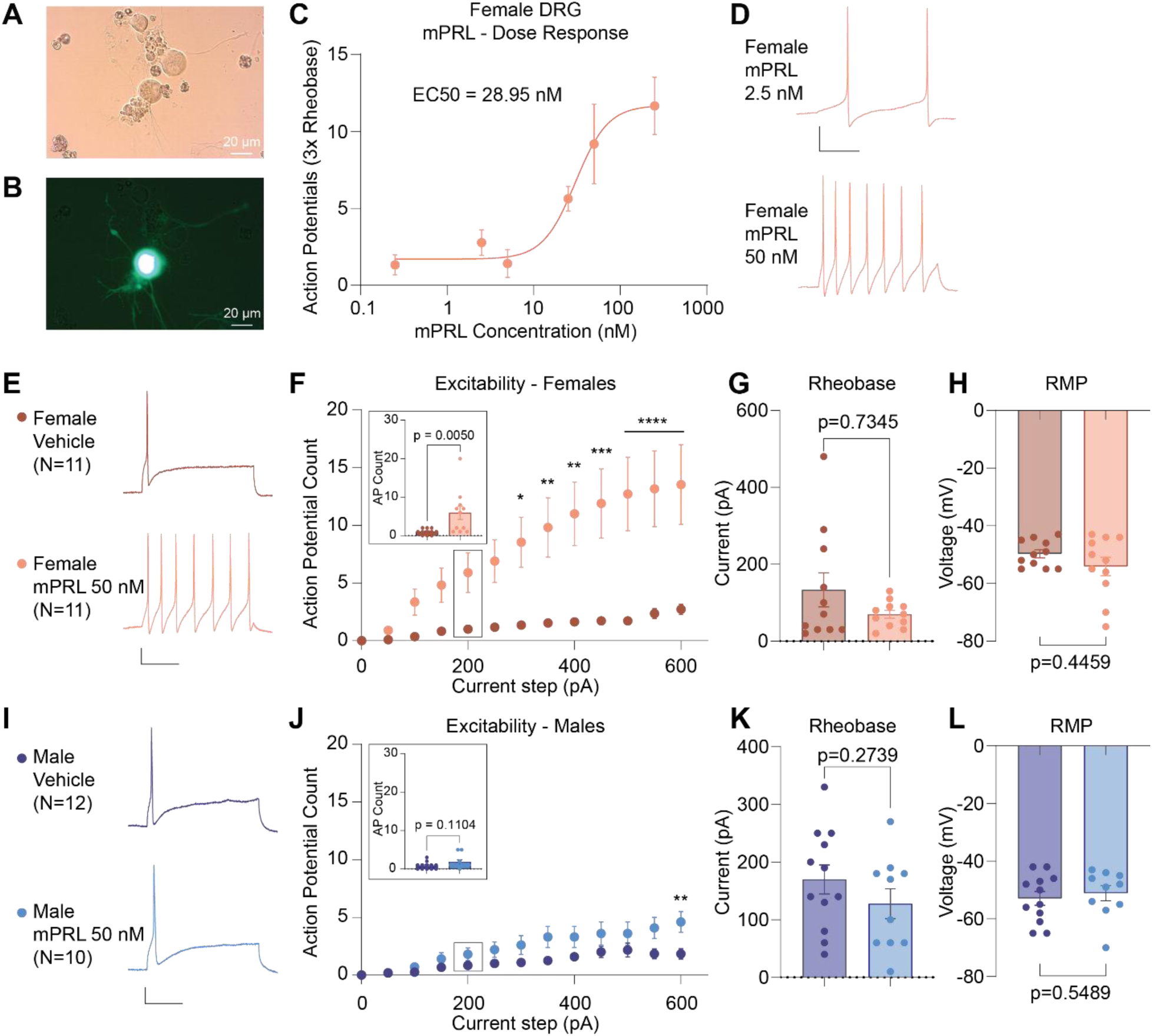
Overnight incubation with mPRL increases excitability of female mouse sensory neurons. **(A)** Representative transmitted light image of cultured lumbar sensory neurons obtained from female Prlr^Cre/+^;Ai6 mice. **(B)** Representative image showing a Prlr-L positive (green) neuron in the culture. **(C)** Concentration response curve using evoked action potential number following a pulse of 3x rheobase demonstrating that the effective concentration of mPRL in vitro is ∼ 29 nM after overnight incubation. **(D)** Representative action potential traces evoked using a depolarizing current pulse equal to 3x the rheobase from female mouse DRG neurons expressing the prolactin receptor after overnight incubation with indicated concentrations of mPRL. **(E)** Representative action potential traces from lumbar DRG neurons isolated from female Prlr^Cre/+^;Ai6 mice showing increased firing after overnight treatment with 50 nM mPRL at the 200-pA current step. **(F)** Excitability of cultured female sensory neurons was increased after overnight incubation with 50 nM mPRL. Inset shows individual data points for the 200-pA current step. **(G)** Rheobase, the minimum current injection to fire a single action potential, was not affected in female DRG neurons after overnight incubation with 50 nM mPRL. **(H)** The resting membrane potential (RMP) was not changed in female DRG neurons after overnight incubation with 50 nM mPRL. **(I)** Representative action potential traces from lumbar DRG neurons isolated from male Prlr^Cre/+^;Ai6 mice showing no change in firing activity after treatment with 50 nM mPRL overnight at the 200-pA current step. **(J)** Excitability of cultured male sensory neurons was not altered after overnight incubation with 50 nM mPRL. Inset shows individual data points for the 200-pA current step. **(K)** The rheobase for male DRG neurons after overnight incubation with 50 nM mPRL was not altered. **(L)** The resting membrane potential for male DRG neurons after overnight incubation with 50 nM mPRL did not change following treatment. Data are represented as mean ± SEM. Differences in excitability were determined with Two-Way ANOVA with Sidak’s multiple comparisons test (F & J). Differences between groups for AP count (F & J inset), rheobase (G & K), and resting membrane potential (H & L) were determined using the Mann-Whitney test *p<0.05, **p<0.01, ***p<0.001, ****p<0.0001. Scale bars represent 20 mV and 100 ms duration. Samples were obtained from at least 3 mice for each condition and the number of cells recorded is indicated in the figure. Details of statistical comparisons can be found in Table S2.

Previous studies suggested that female rodent sensory neurons are selectively sensitized by prolactin for capsaicin-evoked release of calcitonin gene related peptide (CGRP)^14^. We reasoned that if this were true, incubation with mPRL would produce sexually dimorphic hyperexcitability in this mouse neuron system. A robust increase in action potential firing frequency was observed following incubation of female mouse DRG neurons with exogenous mPRL (**Fig. 1E**). This effect was prominent across a range of depolarizing current injection pulses. (**Fig. 1F**). The difference between vehicle and mPRL treated firing patterns is illustrated at the 200-pA current step (**Fig. 1F, inset**). Rheobase, the minimum injected current required to elicit a single action potential, was not affected in cells taken from female mice following overnight treatment with mPRL (**Fig. 1G**) and we did not observe a difference in the resting membrane potential between treatment conditions (**Fig. 1H**). In contrast, the excitability of sensory neurons obtained from male mice was resistant to sensitization by overnight treatment with mPRL (**Fig. 1I**). No difference in excitability was observed in these cells across a range of current steps with only the maximal current injection demonstrating a small, but significant, increase in action potential discharge (**Fig. 1J**). The relative lack of mPRL induced sensitization is illustrated by comparison of the vehicle and mPRL treated male sensory neurons at the 200-pA current step (**Fig. 1J, inset**). We did not observe a difference in rheobase (**Fig. 1K**) or in the resting membrane potential in male mouse DRG (**Fig. 1L**). These results show that mPRL produces robust hyperexcitability in female, but not male, rodent sensory neurons and support the conclusion that PRL sensitization is female selective, but not female specific^23^.

We then explored possible PRL-related sexual dimorphism in human nociceptors. We obtained DRG tissue from consented organ donors and found that the prolactin receptor was robustly expressed in DRG sections recovered from females (**Fig. 2A & Fig. S1A** but that staining was much lower in DRG tissues from male donors (**Fig. 2B & Fig. S1B**). In sensory neurons cultured from female donors, intense cell surface staining for the prolactin receptor in NeuN identified neurons was observed (**Fig 2C**). Much lower intensity signal was observed in cultures prepared from male donors (**Fig. 2D**). These results demonstrate that the prolactin receptor is expressed at much higher levels in female, compared to male, human DRG neurons.

**Figure 2.**
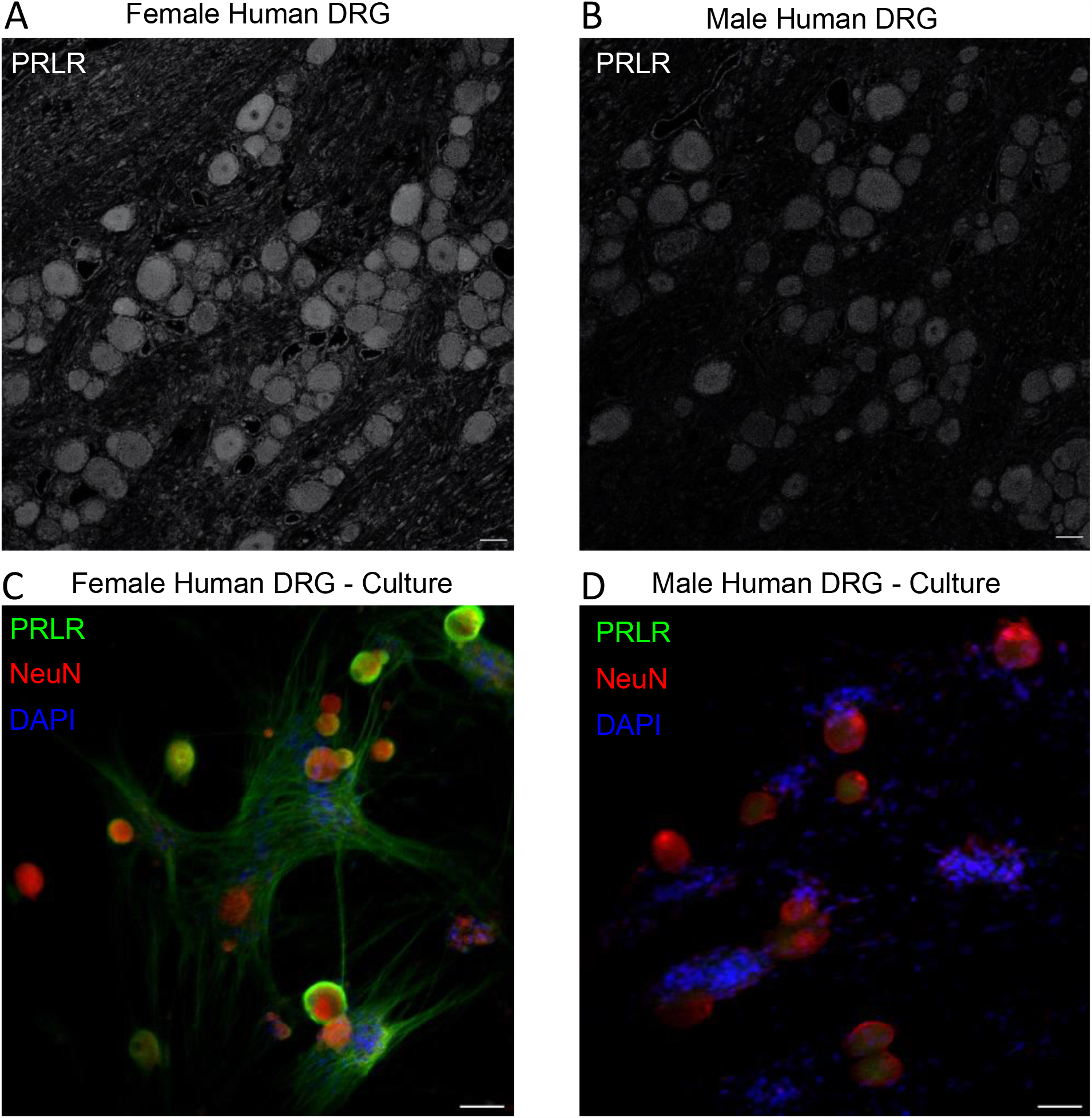
PRLR is expressed at higher levels in female human DRG neurons compared to male DRG neurons. **(A)** Representative immunostaining of DRG tissue from four human female donors illustrating high levels of PRLR expression. **(D)** Male human DRG tissue section (representative of three donors) with markedly lower expression levels of PRLR. **(C)** Representative image of female cultured human DRG stained for PRLR, NeuN, and DAPI, showing robust expression of the receptor on the neuronal membrane. **(D)** Representative image of male human DRG stained for PRLR, NeuN, and DAPI with much lower levels of receptor expression. Scale bars are 50 μm.

We next sought to investigate whether PRL-induced sensitization was sexually dimorphic in human sensory neurons. DRG neurons from female and male human donors were cultured overnight with recombinant human prolactin (hPRL) and patch clamp electrophysiology was performed to determine possible effects on action potentials elicited by increasing stepwise depolarizing current pulses. In female sensory neurons, treatment with 50 nM hPRL induced robust hyperexcitability (**Fig. 3A**) observed across a range of current injection steps (**Fig 3B**). The difference in firing pattern between vehicle and hPRL treated neurons is illustrated at the 1500 pA current step (**Fig 3B, inset**). The rheobase of these neurons was marginally reduced following hPRL treatment (**Fig. 3C**). As in mouse DRG neurons, there was no difference in the resting membrane potential between neurons treated overnight with vehicle and those treated with hPRL (**Fig. 3D**). In contrast, when we incubated sensory neurons from male donors with hPRL we did not observe an increase in excitability between vehicle and hPRL treatment (**Fig. 3E & F, Fig3F inset**). There was also no change in the rheobase (**Fig. 3G**) or the resting membrane potential in male sensory neurons (**Fig. 3H**). These results demonstrate that hPRL sensitizes female human sensory neurons but not male human sensory neurons. It should be noted that we observed mPRL-induced reduced rheobase in rodent, but not human studies. Rheobase is primarily determined by threshold setting channels such as the voltage gated sodium channel Nav1.7 and T-type calcium channels^24, 25^ and it is possible that ion channel composition varies significantly between human and rodent sensory neurons^26, 27^, potentially explaining the species difference observed here.

**Figure 3.**
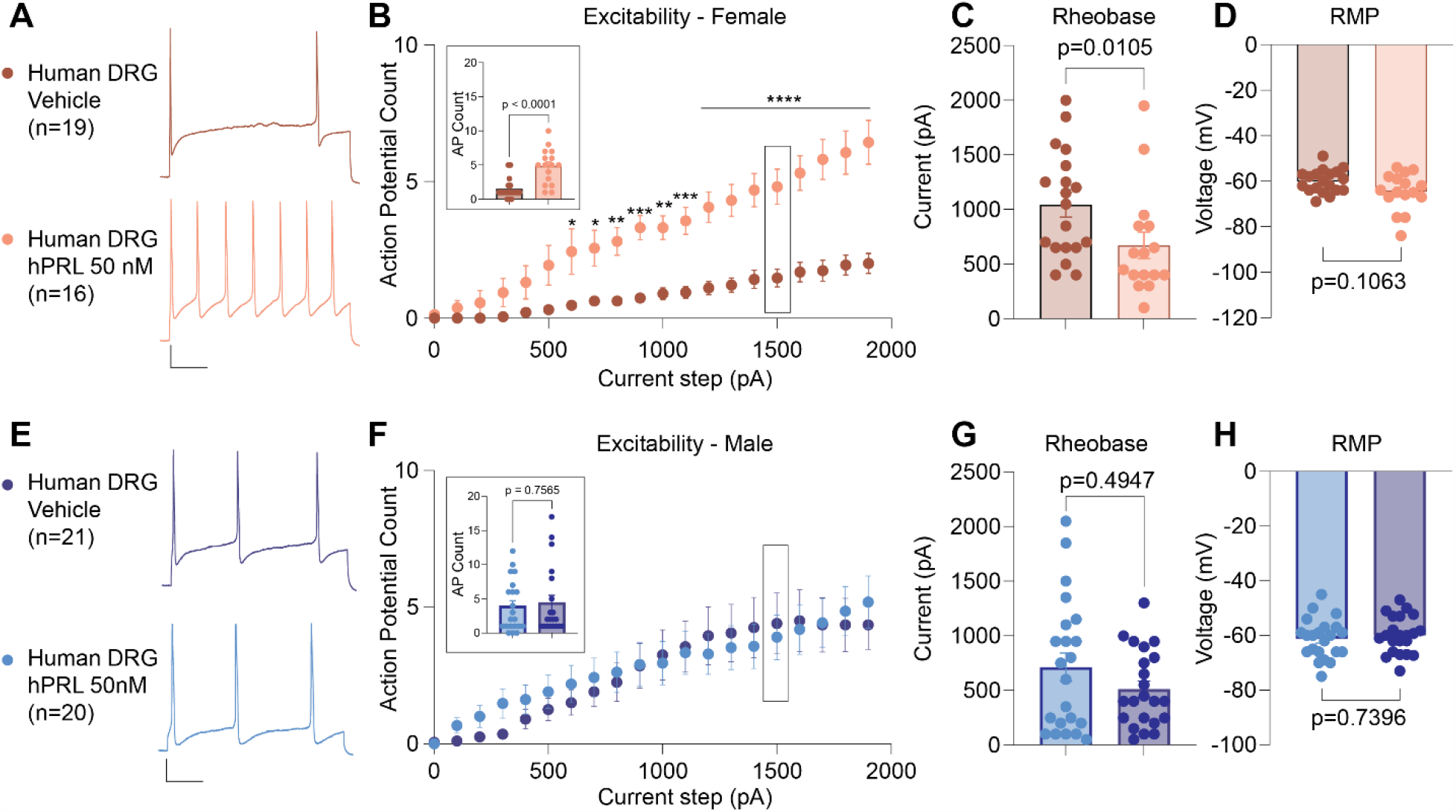
Overnight incubation with hPRL increases excitability of female human sensory neurons. **(A)** Representative action potential traces showing increased firing from female human sensory neurons after overnight incubation with 50 nM hPRL, which were evoked with a depolarizing current pulse to 1500 pA. **(B)** Excitability of cultured human female sensory neurons was increased after overnight incubation with 50 nM hPRL. Inset shows individual data points for the 1500-pA current step. **(C)** Rheobase for human female sensory neurons after overnight incubation with 50 nM hPRL decreased slightly. **(D)** The resting membrane potential (RMP) after overnight incubation with 50 nM hPRL was not changed. **(E)** Representative action potential traces obtained from male human sensory neurons showing no change in firing after overnight incubation with 50 nM hPRL, which were evoked with a depolarizing current pulse to 1500 pA. **(F)** Excitability of cultured human male sensory neurons did not change after overnight incubation with 50 nM hPRL. Inset shows individual data points for the 1500-pA current step. **(G)** Rheobase for human male human sensory neurons after overnight incubation with 50 nM hPRL was unchanged. **(H)** The resting membrane potential for male human sensory neurons after overnight incubation with 50 nM hPRL was unchanged. Data are represented as mean ± SEM. Differences in excitability were determined with Two-Way ANOVA with Sidak’s multiple comparisons test (B & F). Differences between groups for AP count, rheobase (C & G), and resting membrane potential (D & H) were determined using the Mann-Whitney test. *p<0.05, **p<0.01, ***p<0.001, ****p<0.0001. Scale bars 20 mV and 200 ms. These recordings were obtained from 2 female and 3 male donors with the number of cells that were recorded for each group indicated in the figure. Details of statistical comparisons can be found in Table S2.

To our knowledge, these data represent the first demonstration of functional sexual dimorphism in human nociceptors. Sensitization of primary afferent input has been demonstrated to be a driving force for pain in humans^28, 29^. Non-steroidal anti-inflammatory drugs (NSAIDs), likely the most common analgesic class in the world, act by blocking nociceptor sensitization confirming that blockade of nociceptor sensitization is therapeutically meaningful^30^. NSAIDs, however, have severe side-effects including gastrointestinal and kidney toxicity that limit their overall utility. Our data demonstrate that prolactin is profoundly female selective in promoting sensitization of both rodent and human nociceptors. Therefore, these findings support the conclusion that mechanisms promoting excitability are largely conserved across species and suggest that mechanistic investigations of PRL-induced sensitization in rodents have significant translational relevance. Limited availability of human cells has thus far not allowed detailed investigation of reasons for female selectivity of PRL-induced nociceptor sensitization. One possibility however is that prolactin receptor expression is lower in male than in female human nociceptors as observed in our studies. Thus, while sex differences in prolactin receptor transcripts have not been reported in human nociceptors, sexual dimorphism likely occurs at the protein level.

Taken together, sexually dimorphic differences in prolactin receptor expression and PRL-induced functional outcomes support the emerging and remarkable conclusion that nociceptors, the fundamental building blocks of pain, are different in men and women. Pain may thus be produced by different mechanisms in men and in women suggesting the need for consideration of patient sex in pain therapy. Despite higher prevalence of chronic pain in women, female specific therapies have not been identified. Our findings open a path for first in class prolactin targeted therapies that could be tailored for the treatment of pain in women.

## Methods

### Animals

All procedures were approved by the University of Arizona Institutional Animal Care and Use Committee in accordance with the guidelines of the committee for Research and Ethical Issues of the International Association for the Study of Pain. Female and male 8-12-week-old mice were used in this study. Mice were housed 2-5 per cage and were maintained in a climate-controlled room on a 12-hour light/dark cycle with access to food and water ad libitum.

C57BL/6J mice were purchased from Jackson Laboratories. Prlr^Cre/+^ mice were made by Dr. Ulrich Boehm (University of Saarland School of Medicine, Homberg, Germany) and kindly provided by Dr. Armen Akopian (University of Texas Health Science Center at San Antonio, San Antonio, Texas). In these mice, an internal ribosome entry site (IRES) followed by Cre recombinase cDNA is inserted immediately after exon 10 in the prlr gene. Ai6 green fluorescent protein reporter mice (B6.Cg-Gt(ROSA)26Sortm6(CAG-ZsGreen1)Hze/J; Stock No: 007906) were purchased from Jackson Laboratories. We then crossed the Ai6 mice with the Prlr^Cre/+^ mice to obtain a reporter mouse Prlr^Cre/+^;Ai6 that expressed GFP in Prlr positive cells.

### Culture of mouse dorsal root ganglia neurons

Dorsal root ganglia (DRG) were dissected from adult (∼3 months) male and female Prlr^Cre/+^;Ai6 mice employing procedures described previously (23). Briefly, mice were euthanized with an overdose of isoflurane before decapitation. The lumbar spine was isolated, and a laminectomy was performed to visualize the spinal cord and DRG bodies. The L2-5 ganglia were dissected out and placed in cell culture media before being transferred to media containing neutral protease 3.125 mg/mL (Worthington, LS02104) and collagenase type 1 5 mg/mL (Worthington, LS004194). The ganglia were incubated with the enzyme for approximately 45 minutes before mechanical dissociation with a fire polished Pasteur pipette. Neurons were maintained in neurobasal-A cell culture media (Thermo Fisher, #10888022) supplemented with 2% B-27 (Thermo Fisher, #17504044), 1% Penicillin/Streptomycin (Hyclone, #16777-164), 1% GlutaMax (Thermo Fisher, 35050061), 10% fetal bovine serum (Sigma, #F0926-50ML), 25 ng/mL GDNF (VWR, #10788-106), and 100 ng/mL mouse NGF (VRW, #76046-694). Dissociated DRG neurons were plated onto 12 mm poly-D-lysine and laminin-coated glass coverslips and cultured for up to 24-48 h. After 12 hours in culture, cells were incubated with recombinant mouse PRL overnight and electrophysiological recordings were subsequently performed.

### Human DRG culture

All human tissues that were used for the study were obtained by legal consent from organ donors in the US. AnaBios Corporation’s procurement network includes only US based Organ Procurement Organizations and Hospitals. Policies for donor screening and consent are the ones established by the United Network for Organ Sharing (UNOS). Organizations supplying human tissues to AnaBios follow the standards and procedures established by the US Centers for Disease Control (CDC) and are inspected biannually by the DHHS. Distribution of donor medical information is in compliance with HIPAA regulations to protect donor’s privacy. All transfers of donor tissue to AnaBios are fully traceable and periodically reviewed by US Federal authorities. AnaBios generally obtains donor organs/tissues from adults aged 16–60 years old. Donor DRGs from males and females were harvested using AnaBios’ proprietary surgical techniques and tools and were shipped to AnaBios via dedicated couriers. The DRGs were then further dissected in cold proprietary neuroplegic solution to remove all connective tissue and fat. The ganglia were enzymatically digested, and the isolated neurons put in culture in DMEM F-12 (Gemini Bio-Products CAT#: 900–955. Lot# M96R00J) supplemented with Glutamine 2 mM, Horse Serum 10% (Invitrogen #16050–130), hNGF (25 ng/ml) (Cell Signaling Technology #5221LF), GDNF (25 ng/ml) (ProSpec Protein Specialist #CYT-305) and Penicillin/Streptomycin (Thermo Fischer Scientific #15140–122). Cells were then incubated overnight with human PRL (R&D Systems, 682-PL-050)

### Patch clamp electrophysiology

For current-clamp recordings the external solution contained (in millimolar): 145 NaCl, 3 KCl, 2 CaCl2, 1 MgCl_2_, 10 D-Glucose, and 10 HEPES (pH 7.4 adjusted with NaOH, and mOsm/L= 300). The internal solution was composed of (in millimolar): 130 K-Gluconate, 5 NaCl, 5 KCl, 3 Mg-ATP, 0.3 EGTA, 10 HEPES, and 10 D-Glucose (pH 7.3 adjusted with KOH, and mOsm/L= 277). All salts were obtained from Sigma Aldrich. Recordings were performed at room temperature (22–24°C) and a tight seal was formed between the patch pipette and the cell membrane (>1 GΩ). After seal formation, a holding potential of -60 mV was applied before breaking the membrane with brief pulses of negative pressure to gain intracellular access and establish the whole-cell patch clamp configuration. A gentle switch from voltage clamp mode to current clamp mode was then performed and the current injection was set to 0 pA to measure passive membrane properties, such as the resting membrane potential. DRG neurons with a resting membrane potential (RMP) more hyperpolarized than −40 mV, stable baseline recordings, and evoked spikes that overshot 0 mV were used for experiments and analysis. After establishing whole-cell mode and verifying that the RMP was stable the cell was subjected to depolarizing pulses to elicit action potential firing. These were evoked by current injection steps from 0–600 pA with an increment of 50 pA in 300 ms steps. Rheobase, which is the minimum current to fire a single action potential, was measured by injecting current stepwise starting from -10 pA in increments of 10 pA for 50 ms. The current step that elicited an action potential was then recorded as the rheobase for that cell. For human DRG neurons, procedures were the same as for mouse neurons with a few exceptions. The rheobase was determined using current pulses in 50 pA steps and the excitability was determined using current pulses in 100 pA steps, for 1 s, from 0 to 2000 pA. This range was sufficient to saturate the firing of recorded neurons.

Recording pipettes were pulled from standard wall borosilicate glass capillaries (Sutter Instruments) with a horizontal puller (Model P-1000, Sutter Instruments). The input resistance of the pipettes ranged from 2 to 4 MΩ. Recordings were performed from small DRG neurons with capacitance between 10 and 35 pF (∼18-33 μm, mouse). Series resistance under 7 MΩ was deemed acceptable. All experiments had a series resistance compensation between 60-90 %. Signals were acquired using a HEKA EPC10 USB amplifier, filtered at 10 kHz, and digitized at 20 kHz. Analyses were performed using Fitmaster software (HEKA) and Origin 9.0 software (OriginLab).

### Immunostaining of dissociated human DRG

After completion of electrophysiological recordings, human DRG cells cultured on glass coverslips were fixed for immunostaining. The cells were permeabilized with 0.1% TritonX100, blocked with 1% normal goat serum (NGS) for 1 hour and incubated overnight with primary recombinant anti-prolactin receptor antibody [EPR7184(2)] (abcam, ab170935) at 1:50 in 1% BSA, 1.5% NGS/PBSTX. After three washes, the cells were incubated for 1 hour with secondary goat anti-rabbit Alexa488 (ThermoFisher A-32731) at 1:1000 dilution in 1%BSA, 1.5% NGS/PBSTX. After washing, slides were counterstained for 2 hours with NeuN antibody (MAB377A5 Sigma-Aldrich, Alexa Fluor® 555 Conjugate, 1:100 dilution) and DAPI (1:5000 dilution), washed and mounted with ProLong™ Gold Antifade Mountant (Invitrogen P36930). Confocal images were acquired using an Olympus FV1200 microscope (Olympus Life Sciences, Waltham, MA) and a 20x/0.8 or 40x/0.95 UPLXApo objective using excitation beams of 405, 488, and 561 nm. The confocal images of single sections were acquired from male and female samples and processed in the ImageJ software using identical parameters.

### Human DRG preparation and immunostaining

Human dorsal root ganglia (DRG) were obtained from Mid America Transplant as part of a partnership with Dr. Moutal’s lab. All studies involving human tissues have been classified for an IRB exemption #4 at Saint Louis University. The tissues were fixed by immersion in 10% formalin for at least one day. After fixation, the DRGs were trimmed from their connective tissue and roots, cut in half, prior to embedding into paraffin blocks. The paraffin blocks were sectioned in 5 μm slices. Deparaffination is carried out by three xylene washes for 5 minutes each, followed by two 100% alcohol washes for 1 minute each, then two 95% alcohol washes for 1 minute each, and finally by washing in running water for 2 minutes.

For antibody staining, the slides were washed twice with phosphate buffered saline (PBS) for 5 minutes each at room temperature then exposed for 30 minutes to a blocking solution containing 5% donkey serum and 1% BSA in PBS at room temperature. The primary antibody against PRLR (1:100, Cat# ab170935, Abcam) was then incubated in blocking buffer, overnight at 4°C in a humidified chamber. The slides were then washed 3 times, 15 minutes with PBS at room temperature before adding the secondary antibody solution (donkey anti-rabbit, alexa594, 1/300) for one hour at room temperature. Slides were then washed with PBS 3 times for 15 minutes at room temperature. Finally, slides were mounted in ProLong Gold (Cat# P36930, thermo fisher scientific) to protect from fading and photobleaching. Immunofluorescent micrographs were acquired on a Leica SP8 inverted upright microscope using a 10x dry objective. For a clearer and more reliable signal the acquisition window of the detector was set between 602 and 637nm, which corresponds to 15nm above and below the emission peak of Alexa 594. For all quantitative comparisons among cells under differing experimental conditions, camera gain and other relevant settings were kept constant. The freeware image analysis program Image J (http://rsb.info.nih.gov/ij/) was used for extracting representative pictures, no contrast enhancement was performed.

### Statistical methods and data analysis

Graphing and statistical analysis was undertaken with GraphPad Prism (Version 9). All data sets were checked for normality using the D’Agostino & Pearson test. Outliers were identified using the ROUT function with Q set to a value of 1%. Details of statistical tests, significance and sample sizes are reported in the supplementary statistical table. All data plotted represent mean ± SEM. The dose response curve for mPRL in murine DRG neurons was fit using non-linear regression with a variable slope four parameter function. For electrophysiological recordings, excitability was compared across current injection steps and between treatment groups using a Two-way ANOVA with Šídák’s multiple comparisons test. Comparisons made between two groups, including: number of action potential counts at a specific current injection step, rheobase, and the resting membrane potential using a Mann-Whitney test.

## Notes

### Competing Interest Statement

The authors have declared no competing interest.

