## Supplemental Figures and Tables for "Functional Sexual Dimorphism in Human Nociceptors"

**Supplemental Information:**

**Table S1 - Human subjects' demographic information**

| Sex | Age | Ethnicity | Cause of Death |
|---|---|---|---|
| Male | 21 | African American | Anoxia |
| Female | 35 | Caucasian | Head trauma/blunt injury |
| Female | 23 | Caucasian | Anoxia |
| Male | 16 | African American | Head trauma |
| Male | 48 | Hispanic | CVA/ICH/Stroke |
| Female | 25 | Asian | CVA/ICH/Stroke |
| Male | 61 | Caucasian | ICH/Stroke |
| Male | 44 | Caucasian | Anoxia |

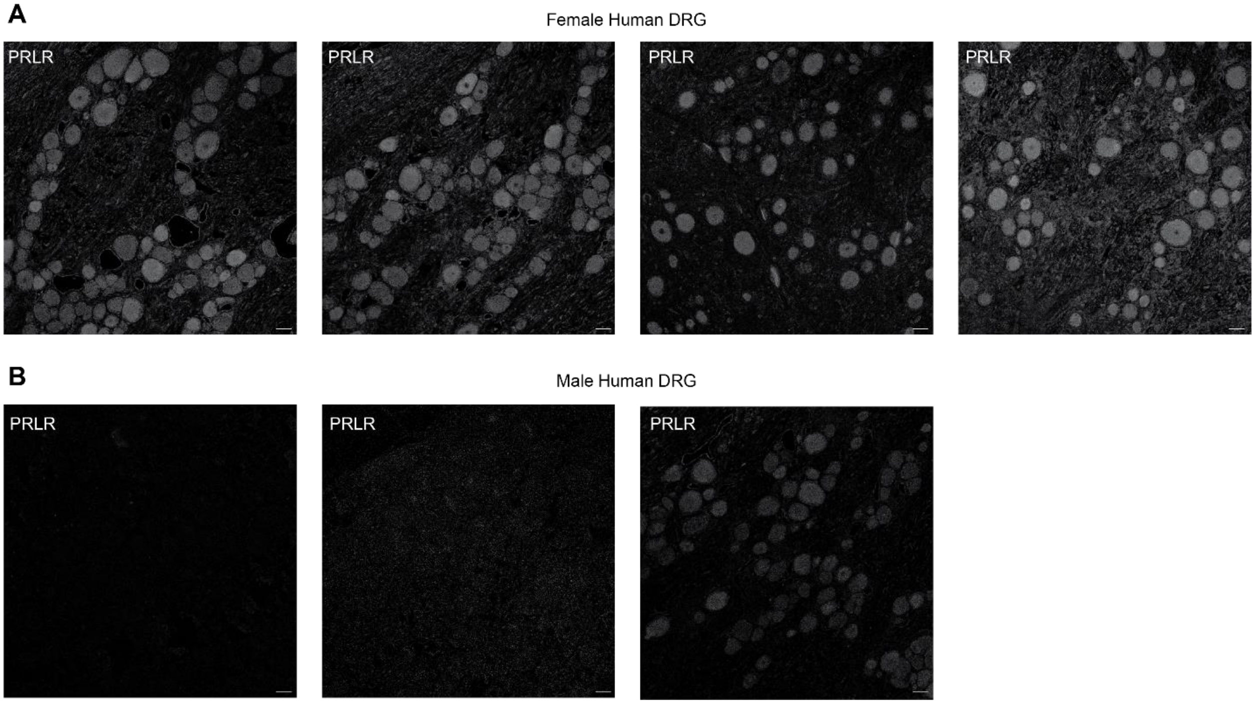


**Figure S1. Immunohistochemical staining for PRLR in female and male human dorsal root ganglia tissue section**. **(A)** Immunostaining showing robust expression of PRLR in DRG obtained from four DRG sections of a human female donor. **(B)** Immunostaining showing that expression of PRLR is nearly absent in DRG tissue from three sections of two male human donors. Scale bars are 50 μm.

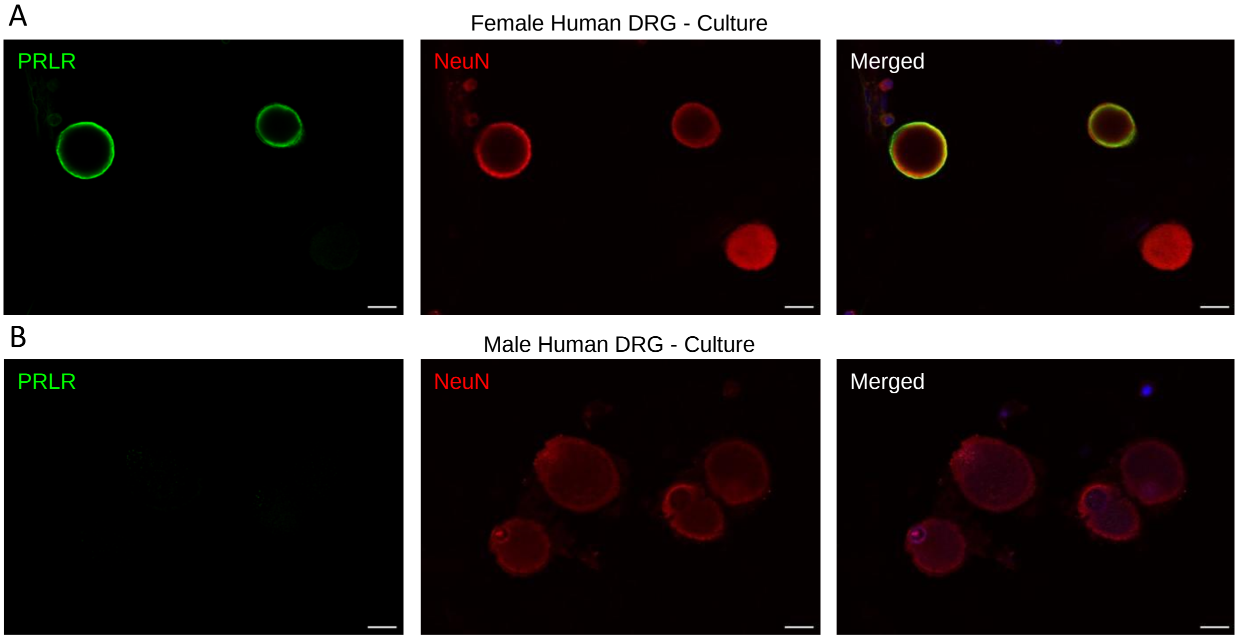


**Figure S2.** Representative confocal images (2.0 µm/slice) of fixed cultured DRG neurons from a female **(A)** and a male **(B)** donor stained for PRLR (left), NeuN (middle), and DAPI shown in the merged image (right). Note the absence of signal for the male PRLR staining compared to the female. Scale bars are 20 µm.

**Table S2. Details of statistical comparisons**

| Figure | Assay | Statistical test and findings | Post hoc test and p values | Number of subjects |
|---|---|---|---|---|
| Figure 1C | mPRL dose response curve | Non-linear regression<br>$R^2 = 0.9073$ | Bottom = 1.741<br>Hillslope = 3.021<br>Top = 11.66<br>EC50 = 28.95 logEC50 = 1.484<br>Span = 10.01<br><br><u>95% Confidence Intervals</u><br>Bottom: 1.316 to 2.162<br>Hillslope: 1.863 to 7.225<br>Top: 10.67 to 12.68<br>EC50: 25.59 to 33.77<br>logEC50: 1.408 to 1.528 | 0.25 nM (n=12)<br><br>2.50 nM (n=9)<br><br>5.0 nM (n=7)<br><br>25 nM (n=11)<br><br>50 nM (n=5)<br><br>250 nM (n=6) |
| Figure 1F | Excitability - female mouse DRG mPRL 50 nM | Two-Way ANOVA<br><br>Interaction: p=0.0019<br><br>F (12, 260) = 2.699 | Sidak's multiple comparisons<br><br>Female vehicle vs. Female mPRL<br><br>P values<br>0 pA p>0.9999<br>50 pA p>0.9999<br>100 pA p= 0.9476<br>150 pA p= 0.7063<br>200 pA p= 0.3932<br>250 pA p= 0.1836<br>300 pA p= 0.0317<br>350 pA p= 0.0066<br>400 pA p= 0.0011<br>450 pA p= 0.0003<br>500 pA p<0.0001<br>550 pA p<0.0001<br>600 pA p<0.0001 | Female vehicle (n=11)<br><br>Female mPRL (n=11) |
| Figure 1F Inset | Female mouse - action potentials fired at 200 pA | Mann-Whitney test | Female vehicle vs. Female mPRL<br>P=0.0050 | Female vehicle (n=11)<br><br>Female mPRL (n=11) |
| Figure 1G | Rheobase - female mouse DRG mPRL 50 nM | Mann-Whitney test | Female vehicle vs. Female mPRL<br>P=0.7345 | Female vehicle (n=11)<br><br>Female mPRL (n=11) |
| Figure 1H | Resting membrane potential - female mouse DRG mPRL 50 nM | Mann-Whitney test | Female vehicle vs. Female mPRL<br>P=0.4459 | Female vehicle (n=11)<br><br>Female mPRL (n=11) |

| | | | | |
|---|---|---|---|---|
| Figure 1J | Excitability -male mouse DRG mPRL 50 nM | Two-Way ANOVA<br><br>Interaction: p=0.3167<br><br>F (12, 260) = 1.154 | Sidak's multiple comparisons<br><br>P values<br>0 pA p>0.9999<br>50 pA p>0.9999<br>100 pA p>0.9999<br>150 pA p= 0.9962<br>200 pA p= 0.9581<br>250 pA p= 0.8244<br>300 pA p= 0.5073<br>350 pA p= 0.1122<br>400 pA p= 0.3149<br>450 pA p = 0.4221<br>500 pA p= 0.5960<br>550 pA p= 0.0506<br>600 pA p= 0.0060 | Male vehicle (n=12)<br><br>Male mPRL (n=10) |
| Figure 1J Inset | Male mouse - action potentials fired at 200 pA | Mann-Whitney test | Male vehicle vs. Male mPRL<br>P=0.1104 | Male vehicle (n=12)<br><br>Male mPRL (n=10) |
| Figure 1K | Rheobase - male mouse DRG mPRL 50 nM | Mann-Whitney test | Male vehicle vs. Male mPRL<br>P=0.2739 | Male vehicle (n=12)<br><br>Male mPRL (n=10) |
| Figure 1L | Resting membrane potential - male mouse DRG mPRL 50 nM | Mann-Whitney test | Male vehicle vs. Male mPRL<br>P=0.5489 | Male vehicle (n=12)<br><br>Male mPRL (n=10) |
| Figure 3 B | Excitability - female human DRG mPRL 50 nM | Two-Way ANOVA<br><br>Interaction: p<0.0001<br><br>F (19, 660) = 4.561 | Sidak's multiple comparisons<br>0 pA p>0.9999<br>100 pA p>0.9999<br>200 pA p=0.9998<br>300 pA p=0.9546<br>400 pA p=0.7579<br>500 pA p=0.1384<br>600 pA p=0.0238<br>700 pA p=0.0286<br>800 pA p=0.0065<br>900 pA p=0.0005<br>1000 pA p=0.0014<br>1100 pA p=0.0003<br>1200 pA p<0.0001<br>1300 pA p<0.0001<br>1400 pA p<0.0001<br>1500 pA p<0.0001<br>1600 pA p<0.0001<br>1700 pA p<0.0001<br>1800 pA p<0.0001<br>1900 pA p<0.0001<br>2000 pA p<0.0001 | Female human vehicle (n=19)<br><br>Female human mPRL (n=16) |

| | | | | |
|---|---|---|---|---|
| Figure 3B Inset | Female human - action potentials fired at 1500 pA | Mann-Whitney test | Female human vehicle vs. Female human hPRL P<0.0001 | Female human vehicle (n=19) Female human mPRL (n=16) |
| Figure 3C | Rheobase - female human DRG mPRL 50 nM | Mann-Whitney test | Female human vehicle vs. Female human hPRL P=0.0105 | Female human vehicle (n=19) Female human mPRL (n=16) |
| Figure 3D | Resting membrane potential - female human DRG mPRL 50 nM | Mann-Whitney test | Female human vehicle vs. Female human hPRL P=0.1063 | Female human vehicle (n=19) Female human mPRL (n=16) |
| Figure 3F | Excitability -male human DRG mPRL 50 nM | Two-Way ANOVA<br><br>Interaction: p=0.9995<br><br>F (20, 819) = 0.2699 | Sidak's multiple comparisons<br><br>Male human vehicle vs. Male human hPRL<br>P<0.0001<br>0 pA p>0.9999<br>100 pA p>0.9999<br>200 pA p>0.9999<br>300 pA p=0.9994<br>400 pA p>0.9999<br>500 pA p>0.9999<br>600 pA p>0.9999<br>700 pA p>0.9999<br>800 pA p>0.9999<br>900 pA p>0.9999<br>1000 pA p>0.9999<br>1100 pA p>0.9999<br>1200 pA p<>0.9999<br>1300 pA p>0.9999<br>1400 pA p>0.9999<br>1500 pA p>0.9999<br>1600 pA p>0.9999<br>1700 pA p>0.9999<br>1800 pA p>0.9999<br>1900 pA p>0.9999<br>2000 pA p>0.9999 | Male human vehicle (n=21)<br><br>Male human hPRL (n=20) |
| Figure 3F Inset | Male human - action potentials fired at 1500 pA | Mann-Whitney test | Male human vehicle vs. Male human hPRL P=0.7565 | Male human vehicle (n=21) Male human hPRL (n=20) |
| Figure 3G | Rheobase - male human DRG mPRL 50 nM | Mann-Whitney test | Male human vehicle vs. Male human hPRL P=0.4947 | Male human vehicle (n=21) Male human hPRL (n=20) |
| Figure 3H | Resting membrane potential - male human DRG mPRL 50 nM | Mann-Whitney test | Male human vehicle vs. Male human hPRL P=0.7396 | Male human vehicle (n=21) Male human hPRL (n=20) |
